# Genomic Discovery of EF-24 Targets Unveils Antitumorigenic Mechanisms in Leukemia Cells

**DOI:** 10.1101/2024.10.14.618197

**Authors:** Ajeet P. Singh, Noah Wax, James Duncan, Ana Fernandes, Jonathan L. Jacobs

## Abstract

Curcumin, a polyphenolic compound derived from the plant *Curcuma longa* L., has demonstrated a wide range of therapeutic properties, including potential anticancer effects. However, its clinical efficacy is limited due to poor bioavailability and stability. To overcome these challenges, curcumin analogs like EF-24 have been developed with improved pharmacological properties. In this study, we used whole-transcriptome profiling to identify the genome-wide functional impacts of EF-24 treatment in leukemia cells to improve our understanding of its potential mechanisms of action. This approach allowed us to establish a model system for associating druggable genes with clinical disease targets. We used the chronic myeloid leukemia (CML) cell line K-562 and acute myeloid leukemia (AML) cell lines HL-60, Kasumi-1, and THP-1 to conduct EF-24 treatment studies. Cell viability was significantly decreased in the EF-24–treated cells as compared to the untreated controls. We discovered that the genes ATF3, CLU, HSPA6, OSGIN1, ZFAND2A, and CXCL8, which are associated with reduced cell viability and proliferation, were consistently upregulated in all EF-24–treated cell lines. Further analysis of the tested cell lines revealed the activation of various signaling pathways, including the STAT1 regulated S100 family signaling pathway, that controlled downstream gene expression in response to EF-24 treatment. Our results elucidate the molecular mechanisms underlying EF-24’s antitumor efficacy against leukemia, highlighting its multifaceted impact on signaling pathways and gene networks that regulate cell survival, proliferation, and immune responses in myeloid leukemia cells.

**Significance:** This study reveals how EF-24, a curcumin analog, disrupts key signaling pathways in chronic and acute myeloid leukemia cell lines, offering insights to enhance targeted therapies and provide new targets for investigation.

**Conflict of Interest Statement:** The authors declare no potential conflicts of interest.

## Introduction

Chronic (CML) and acute myeloid leukemia (AML) are heterogeneous hematological malignancies that arise from the rapid uncontrolled proliferation of immature or abnormal myeloid blood cells.^1^ The complexity of these diseases is further exacerbated by diverse subtypes, each characterized by distinct genetic alterations and clinical behaviors. While traditional chemotherapy has improved survival rates for many leukemia patients, there remains a critical need for more targeted and less toxic therapeutic approaches.^2^

Curcumin, a bioactive polyphenolic compound extracted from the rhizomes of *Curcuma longa* L., has garnered significant attention for its potential anticancer properties.^3^ Despite its promising effects in preclinical studies, curcumin’s clinical translation has been hindered by its poor bioavailability and rapid metabolism. To overcome these limitations, researchers have developed synthetic analogs of curcumin with enhanced pharmacological properties and improved efficacy against various cancer types.^4^ EF-24, a structural derivative of curcumin, has demonstrated potent antiproliferative and pro-apoptotic effects in various cancer cell lines and animal models.^3^ Its ability to modulate multiple signaling pathways involved in cell cycle regulation, apoptosis, and inflammation positions EF-24 as a promising candidate for leukemia therapy.^5,6^ However, the precise mechanisms underlying EF-24’s antileukemic effects and its specific molecular targets in leukemia subtypes remain to be elucidated.

To identify the genome-wide functional targets of EF-24 and understand its underlying mechanisms in leukemia cells, we used the CML cell line K-562 (ATCC^®^ CCL-243™) and AML cell lines HL-60 (ATCC^®^ CCL-240™), Kasumi-1 (ATCC^®^ CRL-2724™), and THP-1 (ATCC^®^ TIB-202™) to conduct EF-24 treatment studies. These cell lines represent specific myeloid leukemia subtypes and have been extensively characterized and widely used in research, offering unique insights into the pathogenesis and therapeutic vulnerabilities of the disease. Here, our objective was to first illuminate the baseline transcriptional profiles of these leukemia cell lines, emphasizing the existing molecular discrepancies among them, then conduct next-generation sequencing (NGS) to explore global gene expression responses to EF-24 exposure. Through a comparative differential gene expression analysis of untreated and EF-24–treated cells, we uncovered the molecular pathways and gene regulatory networks underlying EF-24’s anticancer effects. Overall, this approach enabled the visualization of global gene expression landscape and the assessment of target gene relative abundance, enabling the identification of molecular discrepancies among cell lines and facilitating the exploration of EF-24 treatment responses in different cell lines and its effects on target genes.

## Results

### EF-24 exhibits potent cell killing activity in leukemia cell lines

We explored the antitumorigenic potential of EF-24 among the leukemia cell lines K-562, Kasumi-1, THP-1, and HL-60; these cell lines were selected as they carry distinct genetic mutations and represent specific disease subtypes. After twenty-four hours following EF-24 treatment, we used the CellTiter-Glo^®^ assay (Promega^®^) to evaluate the viability of treated cells as compared to untreated counterparts and found significant reductions of up to 60% in K-562, 75% in THP-1 and HL-60, and 90% in Kasumi-1 (Figure 1A). Cell counts further confirmed the observed reduction in cell viability, exhibiting a diminished number of viable cells across all four treated cell lines as compared to their untreated counterparts (Figure 1B, 1C). These results not only corroborate previous reports but also underscore the effectiveness of EF-24 in killing of cancer cells and demonstrate promising outcomes against a spectrum of myeloid leukemia disease subtypes.^5,6^

**Figure 1:**
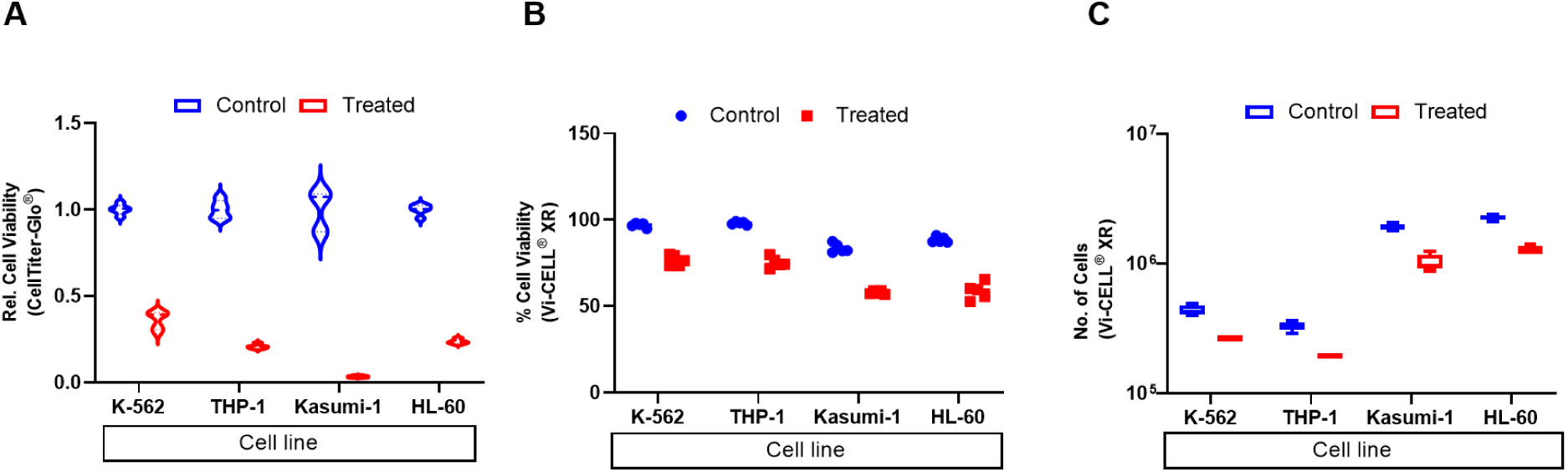
Cell viability in leukemia cell lines is reduced after treatment with EF-24. **(A)** Relative cell viability in control and EF-24 treatment groups as observed by the Promega^®^ CellTiter-Glo^®^ luminescent cell viability assay. **(B)** Percent cell viability and **(C)** the number of cells observed in control and EF-24 treatment groups as observed by the Beckman Coulter^®^ Vi-CELL^®^ XR cell viability analyzer.

### Chronic (CML) and Acute Myeloid Leukemia (AML) cell lines exhibit transcriptional heterogeneity

To understand the inherent molecular characteristics of the myeloid leukemia cell lines, we carried out whole-transcriptome profiling to establish a reference baseline for gene expression in naive cells. Principal component analysis (PCA) revealed close clustering of biological replicates, while leukemia cell lines were stratified into four distinct clusters based on their transcript and gene expression levels, indicating that each of these cell lines represents a distinct model system and transcriptional “starting point” for studying the impact of drug (EF-24) treatment response. Comparative gene expression analysis between the cell lines revealed differential patterns in the top 100 genes, providing insights into their roles in specific cell types (Figure 2A). This facilitated the identification of marker genes specific to each leukemia cell line. For example, *HBG1*, *RHAG*, *HBG2*, *HBA1*, and *NMU* are enriched in K-562 cells as compared to THP-1, Kasumi-1, and HL-60. In contrast, *SMAP2*, *ZMPSTE24*, *CDC20*, *NFYC*, and *RPS4Y1* are enriched in THP-1. Similarly, *CD34*, *KIT*, *EEF1A1*, *PRSS57*, *RPLP0*, and *RPS24* are enriched in Kasumi-1, while *MT-ND4L*, *MT-ND4*, *MT-CO1*, *MT-ND3*, and *CST7* are enriched in HL-60 cells (Figure 2A). The expression patterns of these genes in particular cell lines could serve as biomarkers for characterizing specific leukemia subtypes and as potential pharmacological targets.

**Figure 2:**
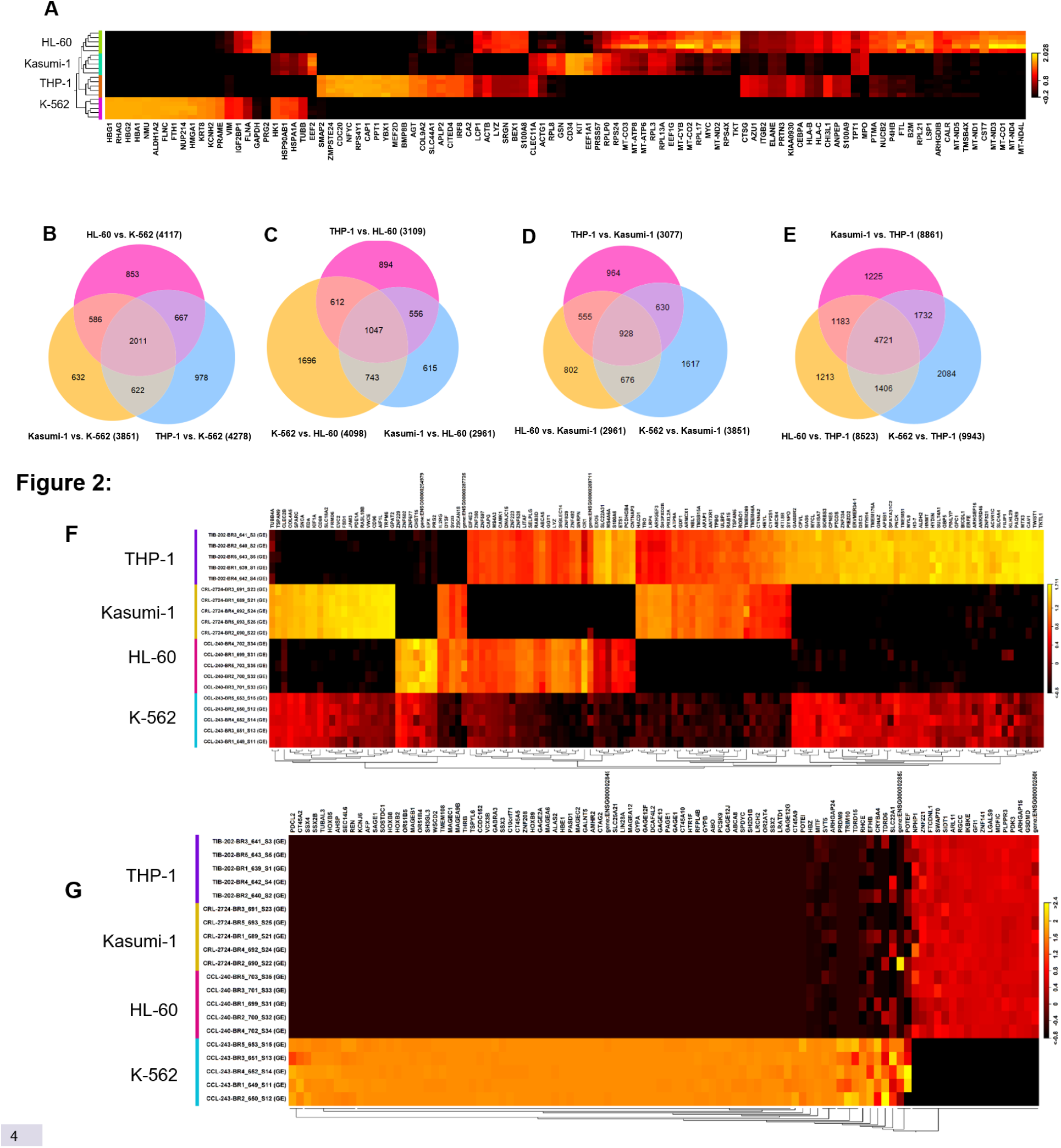
Expression pattern of the top 100 differentially expressed genes that exhibited the highest fold change in chronic myeloid leukemia (K-562) and acute myeloid leukemia (Kasumi-1, HL-60, and THP-1) cell lines. **(A)** Heatmap demonstrating the clustering pattern of genes differentially expressed among leukemia cell lines (red = induced; black = reduced). Each cell line has 5 biological replicates. **(B)** Venn diagram showing the shared and unique genes differentially expressed in the AML subtypes HL-60, THP-1, and Kasumi-1 as compared to the CML subtype K-562. **(C)** Venn diagram showing the shared and uniquely expressed genes in K-562, THP-1, and Kasumi-1 as compared to HL-60. **(D)** Venn diagram showing the shared and uniquely expressed genes in THP-1, K-562, and HL-60 that differ from those in Kasumi-1. **(E)** Venn diagram showing the shared and uniquely expressed genes in Kasumi-1, HL-60, and K-562 that differ from those in THP-1. For each Venn diagram, the differences in circle size reflect the number of genes. **(F)** The heatmap displays the DEGs genes exhibiting inconsistent expression patterns in THP-1, Kasumi-1, and HL-60 as compared to K-562. **(G)** The heatmap displays the genes exhibiting a consistent expression pattern in the THP-1, Kasumi-1, and HL-60 as compared to K-562. Absolute fold change >5, false discovery rate (FDR) p-value 0.01

When further analyzing the number of differentially expressed genes (DEGs) either shared or unique in the AML cell lines as compared to the CML cell line, we identified 2011 genes that were found to be differentially regulated and shared across the THP-1, Kasumi-1, and HL-60 cell lines and distinct from the K-562 cell line (Figure 2B). We also discovered numerous genes unique to HL-60 (853), THP-1 (978), and Kasumi-1 (632) as compared to K-562 (Figure 2B, 2F). Further investigation of HL-60, Kasumi-1, and THP-1 revealed genes that shared similar expression patterns among multiple cell lines as well as those unique to specific cell lines (Figure 2C, 2D, 2E, 2F). We then looked for the genes shared by THP-1, Kasumi-1, and HL-60 that could be a common factor driving the AML subtypes; here, we identified genes with consistent expression patterns among the cell lines. For instance, *PDCL2*, *CT45A2*, *SSX4*, *SSX2B*, *TUBAL3*, *HOXB5*, *AHSP*, and *SEC14L6* are consistently downregulated whereas *GSDMD*, *ARHGAP15*, *PDK3*, *PLPPR3*, *MDFIC*, *LGALS9*, *ZNF141*, and *GFI1* are upregulated in HL-60, Kasumi-1, and THP-1 as compared to K-562 (Figure 2G). Next, we identified genes inconsistently regulated in THP-1, Kasumi-1, and HL-60 cells and differentially expressed as compared to K-562 cells. For example, *TUBB4A4*, *TSPAN9*, *CLEC2B*, *COL4A5*, *SPARC*, and *SNCA* are highly augmented in Kasumi-1 cells but attenuated in HL-60 and THP-1 cells as compared to K-562 cells (Figure 2F). Similarly, we also identified a subset of genes differentially enriched or decreased in HL-60 (e.g., *TUBB4A*, *TSPAN9*, *CLEC2B*, *COL4A5*, *SPARC*, *SNCA* decreased; *ZNF229*, *ZNF502*, *ZNF67*, *CHST15*, *EPX* increased) and THP-1 (e.g., *IL2RG*, *DYSF*, *SV2B*, *ZSCAN18* decreased; *TKTL1*, *TWIST1*, *CAV1*, *MTX3*, *PAQR9*, *KLHL29*, *FILIP1*, *SLC4A4*, *ACVR1C* increased) as compared to K-562 (Figure 2F). This comprehensive analysis revealed the global molecular heterogeneity between cell lines and marker genes, underscoring the importance of evaluating different subtypes when reviewing a new treatment as it may affect each cell line differently.

### EF-24 treatment impacts the expression of genes controlling cell survival and proliferation

To identify the gene expression changes responsible for EF-24’s anticancer effects in leukemia cell lines, we performed RNAseq on EF-24-treated cells and compared them against untreated baseline controls. Principal Component Analysis (PCA) was carried out RNAseq profiles for EF-24 treated and untreated samples, with all biological replicates of each condition clustering together (Figure 3A). Furthermore, differential gene expression analysis highlighted a shift in the pattern of gene expression in EF-24–treated cells as compared to control cells (Figure 3B).

**Figure 3:**
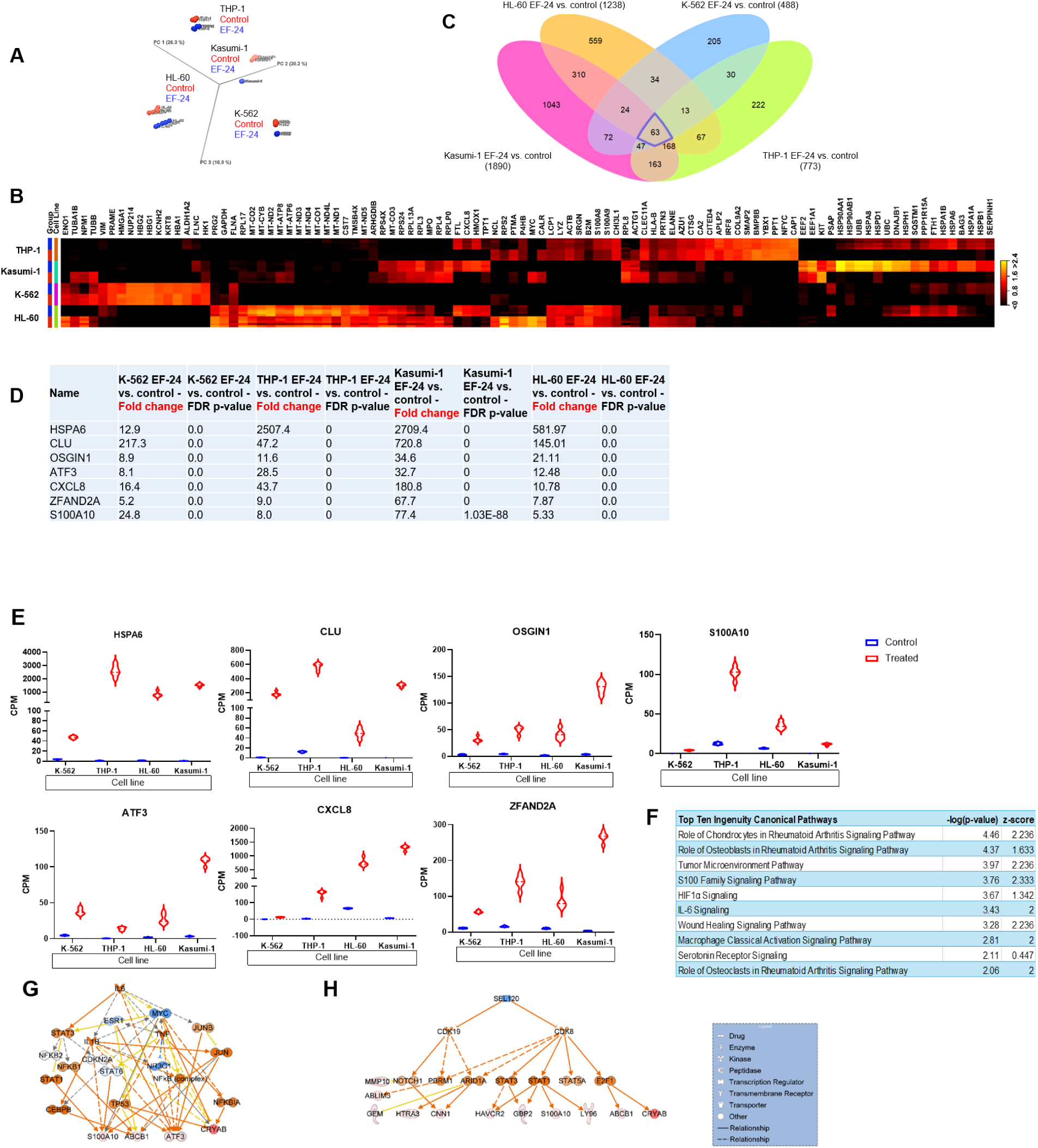
EF-24 treatment causes alterations in key pathways and genes that negatively impact cell survival, proliferation, and viability. **(A)** PCA plot illustrating sample clustering in leukemia cell lines and experimental groups. The variations within the samples resulted in distinct clustering (red = control samples; black = EF-24 samples). **(B)** Heatmap of the top 100 DEGs in leukemia cell lines that shows the highest fold change in EF-24–treated cells as compared to untreated controls. There are 5 biological replicates of each condition/cell line (red = control; blue = treated). **(C)** Venn diagram depicts shared and unique genes that are differentially expressed in EF-24–treated cells as compared to untreated controls. Threshold absolute fold change ≥5, FDR p-value 0.01. **(D)** Pan differentially expressed genes in EF-24–treated cells as compared to untreated controls. (**E**) Violin plots display the relevant genes’ quantitative enrichment in EF-24–treated cells as compared to untreated controls. **(F)** IPA shows shared top canonical pathways activated in myeloid leukemia cell lines were treated with EF-24. **(G)** IL-6 is an upstream regulator of genes induced in the EF-24 cell lines. **(H)** Regulatory network of genes induced in the EF-24 treated cell lines.

Regardless of the distinct myeloid leukemia subtypes, subsequent analysis uncovered 63 differentially expressed genes shared among all four leukemia cell lines treated with EF-24 as compared to the untreated controls of each cell line (Figure 3C). The number of genes altered in individual cell lines following EF-24 treatment were identified (Figure 3C). Several genes known to inhibit cell viability (e.g., *HSPA6*, *CLU*, *OSGIN1*, *ATF3*, *CXCL8*, *ZFAND2A*, and *S100A10*) exhibited a multifold increase in expression level in all four EF-24–treated cell lines as compared to the respective untreated control for each cell line (Figure 3D, 3E). Further, Ingenuity Pathway Analysis (IPA) of genes that were both differentially expressed and shared in all four EF-24–treated leukemia cell lines revealed the top canonical pathways, including the S100 family signaling pathway (Figure 3F). The activation of S100 signaling causes the induction of downstream effector molecules such as *JAK1*, *TYK2*, and *P38 MAPK*, which induce NFkB-regulated proinflammatory cytokines linked to cell proliferation, survival, and viability. ^7^

We then investigated upstream regulators of genes differentially expressed in EF-24-treated cells. IL-6 emerged as a key regulator, activating *JUN*, *NFkB*, and *STAT1* that causing induction of downstream genes S100A10, ABCB1, ATF3 and CRYAB adversely impacting cell viability and survival (Figure 3G). ^8^ Moreover, NUPR1, appeared to be a novel a transcription regulator, upregulated in response to EF-24 treatment and directly control expression of induced genes such as *ATF3, CXCL8, DUSP8, MMP10, UPP1, ZFAND2A,* AND *ADM.* Additionally, networks of upregulated genes in EF-24–treated cells suggest associations with cell death and survival (Figure 3H). Overall, EF-24 treatment causes adverse effects on myeloid leukemia cell lines through alterations in key pathways and genes that negatively impact cell survival, proliferation, and viability. To further understand the transcriptional response to EF-24 therapy in specific leukemia subtypes, we then explored differentially expressed genes and pathways at the individual cell line level.

### EF-24 treatment activated wound healing signaling pathway in K-562 cells

To identify the specific genes and pathways in K-562 cells that are affected by EF-24, we evaluated the genes that were differentially expressed in treated cells as compared to untreated control cells (Figure 4A). Among the differentially regulated genes, 488 genes exhibited a ≥5-fold change (p-value 0.01) in EF-24– treated cells as compared to untreated K-562 controls; only a small subset of DEGs (9 out of 488 genes) were downregulated. We then focused on 25 upregulated genes that showed the highest fold change in the list of DEGs (Figure 4B). Interestingly, *CLU*, *PTPRN*, *NDRG1*, *GBP2*, *OSGIN1*, *ATF3*, *IFI16*, *HLA-C*, *BEX2*, and *VWA5A*—known for their tumor suppressive function—were induced many folds (Figure 4C).^9^

**Figure 4:**
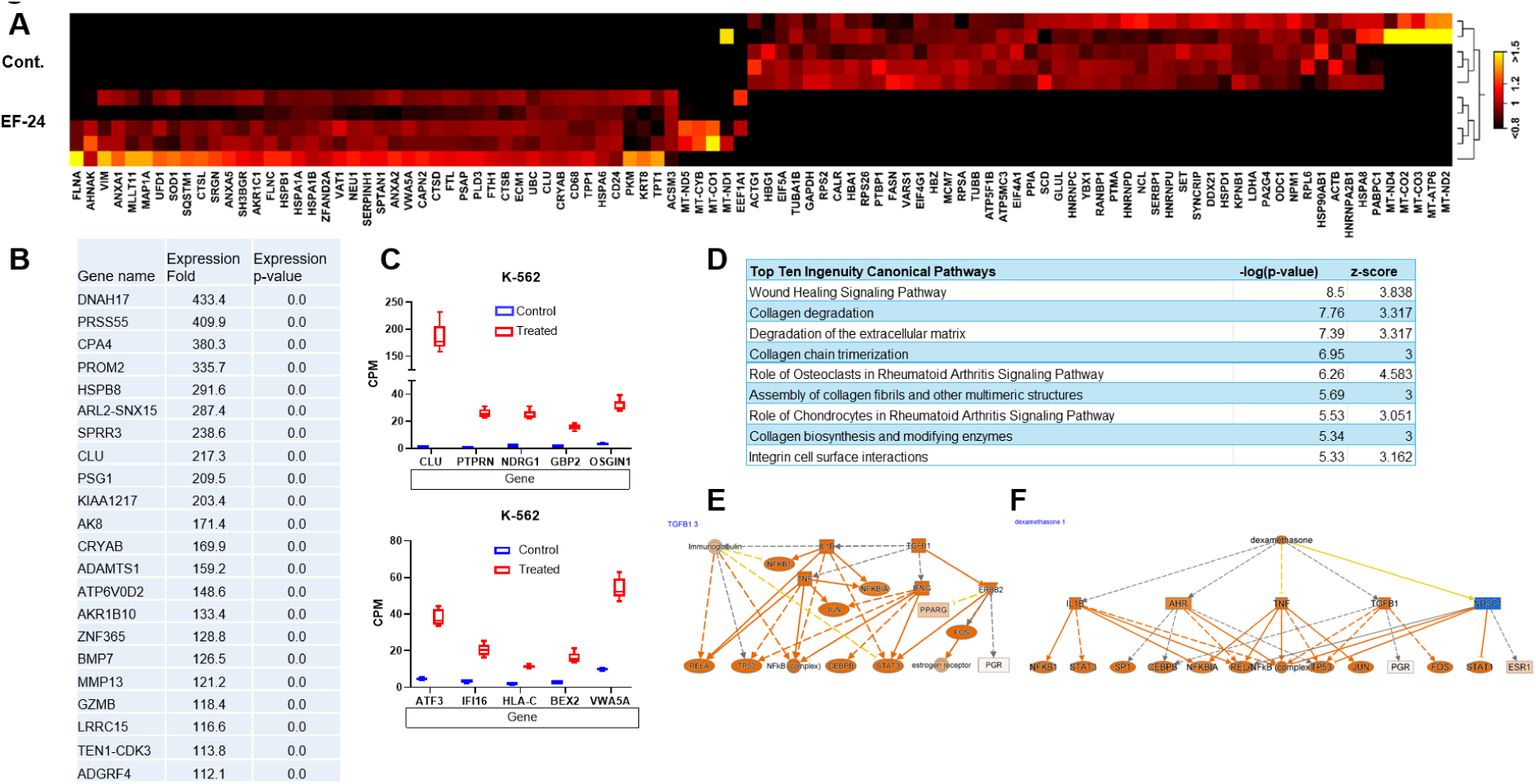
Expression pattern of genes in EF-24–treated K-562 cells and the untreated control. **(A)** Heatmap of the top 100 genes differentially regulated in EF-24–treated cells versus the untreated control (red = induced; black = reduced). Cutoff fold change 5 ≥ p-value 0.01. **(B)** Top 25 molecules changed in EF-24– treated cells versus untreated control K-562 cells. **(C)** Box plots show relative mRNA expression level of indicated genes in EF-24–treated and untreated control cells. ≥5, FDR p-value 0.01. **(D)** IPA shows shared top canonical pathways activated in K-562 cells that were treated with EF-24. **(E, F)** Network of genes predicted to be induced by IL1B and Dexamethasone in K-562 cells treated with EF-24.

Next, analysis in IPA unveiled canonical pathways and gene regulatory networks of the genes activated in response to EF-24 treatment, shedding light on mechanisms influencing cell viability. In this pursuit, IPA illuminated the wound healing signaling pathways and their downstream effector molecules orchestrating the gene expression changes in EF-24–treated cells (Figure 4D). This activation triggered MAP3K7-mediated NFkB and its target genes, which are crucial players in controlling cell survival and death. Additionally, within the wound healing signaling pathway, the EGFR signaling cascade was activated, leading to the activation of downstream genes like *RAS*, *MEK*, and *ERK1/2*. These, in turn, transactivated *AP1* transcription factor–regulated genes that contribute to tissue repair.

Advancing in our transcriptional analysis, we uncovered upstream molecules governing gene expression in EF-24–treated cells (Figure 4E, 4F). The activation of proinflammatory cytokines IL-1B and TGFB1 modulates downstream genes such as *NFKB1*, *RELA*, *TNF*, *JUN*, *TP53*, *CEBPB*, and *FOS*; each of these functions as tumor suppressors (Figure 4E). Meanwhile, the induced expression of various genes, including *STAT3*, *SP1*, and TNF is indirectly promoted and impeded by anti-inflammatory dexamethasone respectively (Figure 4F), which directly activates the repressed NR3C1 glucocorticoid receptor (GR). ^10^ This comprehensive analysis not only enhances our understanding of the molecular responses triggered by EF-24 but also underscores the intricate interplay of pathways and regulatory networks contributing to the observed cellular effects in K-562 cells.

### EF-24–activated pathways cause neutrophil degranulation in HL-60 cells

We performed full transcriptome analysis on HL-60 cells treated or untreated with EF-24 to understand transcriptional alterations and uncover signaling pathways and gene regulatory networks. Differential expression analysis revealed that EF-24 treatment had dramatically changed genes as compared to that of the controls (Figure 5A). Quantitative analysis of DEGs showed that EF-24 treatment significantly increased gene expression levels as compared to untreated controls.

**Figure 5:**
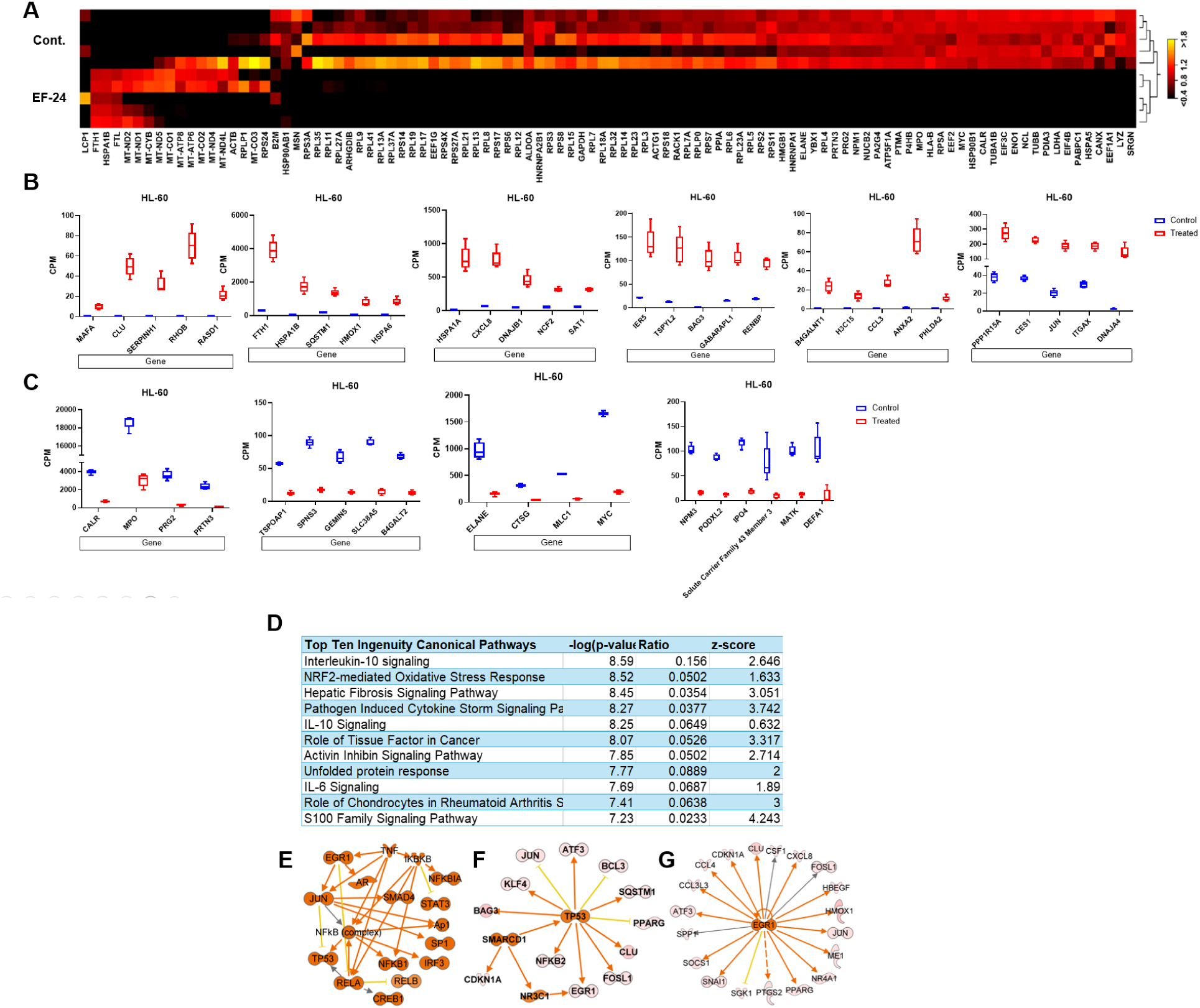
Expression pattern of genes differentially regulated in HL-60 cells treated with EF-24 as compared to the untreated control. **(A)** Heatmap of the top 100 genes altered in HL-60 treated cells as compared to untreated control (red = induced; black = reduced). **(B)** Box plots show relative mRNA level of the indicated genes induced in EF-24–treated as compared to untreated control HL-60 cells. **(C)** Box plots show relative mRNA level of the indicated genes reduced in EF-24–treated as compared to untreated control HL-60 cells. Threshold absolute fold change ≥5, FDR p-value 0.01. **(D)** IPA shows top canonical pathways activated in HL-60 cells were treated with EF-24. **(E)** Gene network of genes regulated by TNF in HL-60 cells treated with EF-24.**(F, G)** Causal gene networks controlled by SMARCD1 and EGR1 in EF-24–treated HL-60 cells.

To understand the functional implications of top DEGs, we employed gene ontology analysis with GeneCards. The analysis indicated that the genes *MAFA*, *CLU*, *RHOB*, *RASD1*, *HSPA1B*, *HSPA1A*, *HMOX1*, *CXCL8*, *DNAJB1*, *NCF2*, *SAT1*, *IER5*, *BAG3*, *B4GALNT1*, *CCL3*, *ANXA2*, *PPP1R15A*, *CES1*, and *JUN*, which were induced in EF-24–treated cells (Figure 5B), may participate in cancer progression.^11,9,12,13^ However, it is noteworthy that their excessive expression level causes adverse effects such as the disruption of homeostasis and cell death.^12,14,15^ This indicates that the overexpression of genes that promote tumor growth affected cellular homeostasis, resulting in a decrease in cell survival and an increase in cell death. Conversely, the genes *FTH1*, *SQSTM1*, *HSPA6*, *TSPYL2*, *PHLDA2*, *ITGAX*, and *DNAJA4* are expected to act as versatile protein regulators, potentially suppressing tumor cell genetic instability and promoting autophagy, apoptosis, and other forms of cell death.^16–21^

We then focused on down-regulated genes associated with cancer, revealing several genes that play pivotal roles in various aspects of cancer development, including cell proliferation, adhesion, migration, phagocytosis, integrin signal transduction, and immunogenic cell death (ICD) (Figure 5C). Interestingly, certain repressed genes like *B4GALT2*, *ELANE*, *CTSG*, *MLC1*, *NMP3*, and *SLC43A3* act as cancer suppressors, inhibiting malignant features and cancer development.^22–24^ This suggests that EF-24 triggers a defense process under stressed physiological conditions that influences both protumorigenic and antitumorigenic gene expression.

Using IPA, we explored the pathways and upstream regulators of DEGs in EF-24–treated HL-60 cells and identified a downregulation of the neutrophil degranulation pathway. This downregulation contributes to an unfavorable environment for leukemia cell survival, leading to cell death.^25^ On the other hand, several signaling pathways, with IL-10 ranking as the top upregulated pathway, were identified (Figure 5D). IL-10 activation was associated with the upregulation of genes such as *HMOX1*, *DUSP1*, *CDKN1a*, *JUN*, *NFKB2*, *BLVRB*, *IL1R1*, *IL1RN*, *BCL3*, and *RELB*, which have all been linked to cell death.^26^ Considering the dual role of IL-10 in cancer biology, with both tumor-promoting and tumor-suppressive effects reported, it suggests that IL-10 activation induced by EF-24 treatment contributes to cell death in HL-60 cells.

Notably, EF-24 treatment activated signaling cascades, including STAT3, that traditionally impede apoptosis, support cell proliferation, facilitate angiogenesis, and suppress antitumor immune responses.^27^ This paradoxical activation of proinflammatory and cell survival pathways suggests a unique mechanism of EF-24-induced cell death in leukemia subtypes. TNF and dexamethasone modulated the expression of altered genes in EF-24 treated HL-60 cells and function as upstream regulator (Figures 5E). Causal network analysis in IPA highlighted SMARCD1 as an upstream regulator, playing a vital role in gene expression regulation through chromatin remodeling.^28^ SMARCD1 activity directly transactivates *P53*, CDKN1a, and *BG2* in EF-24–treated HL-60 cells (Figure 5F), inhibiting cell viability. Additionally, EGR1 activation and its target genes were observed in EF-24–treated cells (Figure 5G). Functional annotation revealed that the genes were activated associated with activation of leukocytes, differentiation of hematopoietic progenitor cells, and cell death of tumor. In summary, our comprehensive analysis of EF-24–treated HL-60 cells revealed a complex interplay of gene expression changes, signaling pathways, and regulatory networks contributing to cell death. The findings underscore the potential of EF-24 as a promising agent for leukemia treatment, warranting further investigation in preclinical and clinical settings.

### EF-24 activates tumor suppressor genes in Kasumi-1 cells

To investigate the gene expression patterns in Kasumi-1 cells in response to EF-24, we conducted a differential gene expression analysis in treated versus untreated cells. Our findings revealed DEGs were either upregulated or downregulated in EF-24–treated cells when compared to control cells (Figure 6A). The treatment with EF-24 induced a significant multifold change in the relative expression levels of various genes; notably, many of these induced genes, including *FTL*, *UBC*, *FTH1*, *UBB*, *SQSTM1*, *JUN*, *MLLT1*, *GADD45A*, *KLF6*, *PPP1R15A*, *TSPYL2*, *SRGN*, *SRXN1*, *NDRG1*, and *ZFP36*, function as tumor suppressors. In contrast, tumor promoters such as *MYBL2*, *TYMS*, *MYC*, *MCM7*, and *EGFL7* were downregulated in EF-24–treated cells (Figure 6B).

**Figure 6:**
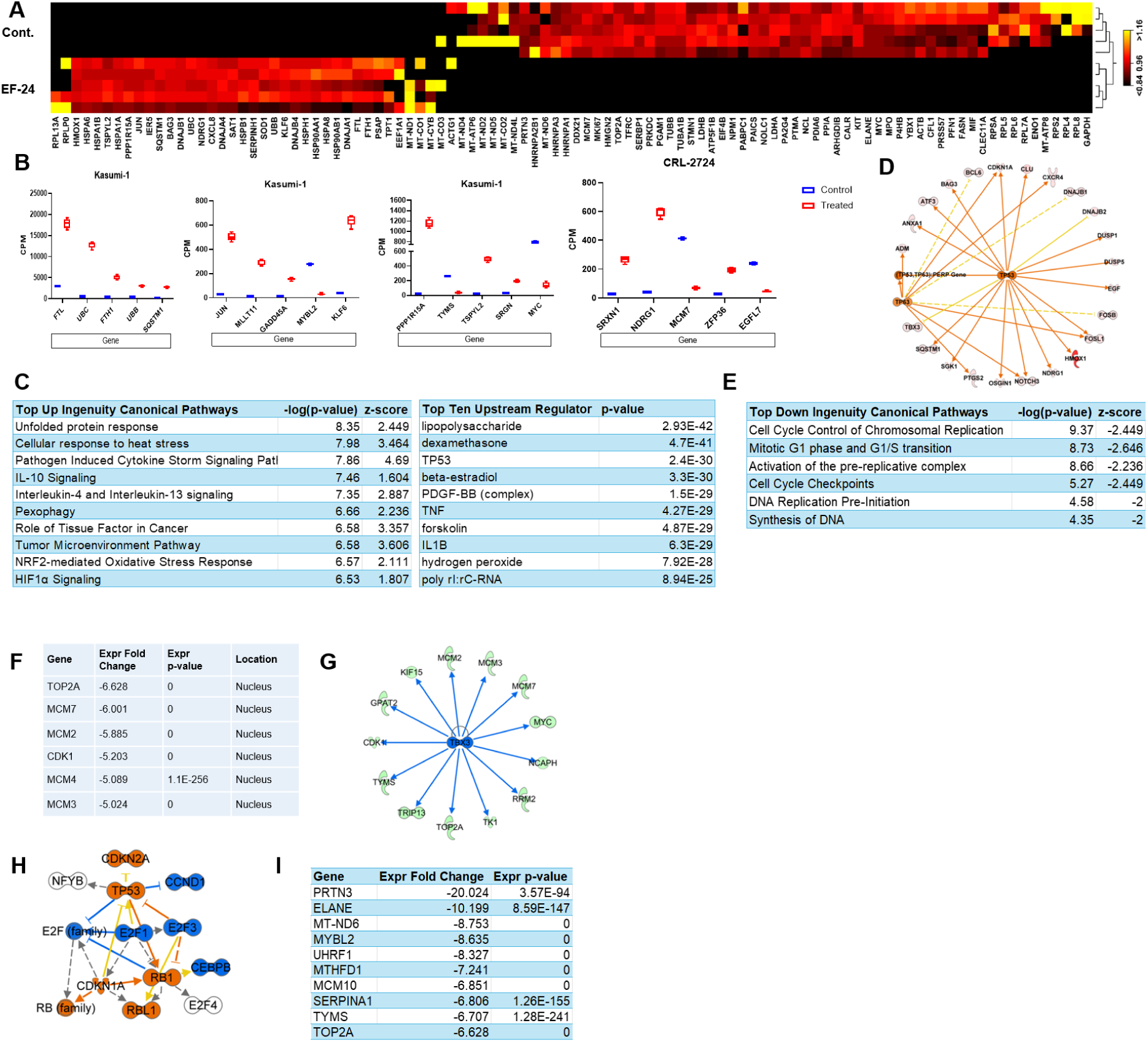
Gene expression changes in Kasumi-1 cells treated with EF-24 as compared with the untreated control. **(A)** Heatmap of the top 100 genes differentially expressed in the EF-24–treated Kasumi-1 cells (red = induced; black = reduced). Absolute fold change >5, FDR p-value 0.01. **(B)** Box plots display the relative levels of candidate genes differentially expressed in the RNA-Seq datasets of EF-24–treated Kasumi-1 cells. **(C)** Top canonical pathways activated in EF-24–treated Kasumi-1 cells. **(D)** TP53 and TP63 are the upstream regulators of genes induced in the EF-24–treated Kasumi-1 cells. **(E)** Top canonical pathway down regulated in the EF-24–treated Kasumi-1 cells. **(F)** Cell cycle and chromosomal replication pathway associated with down regulated genes in the EF-24–treated Kasumi-1 cells. **(G, H)** Network of down regulated genes in the EF-24–treated Kasumi-1 cells regulated by TBX3 and E2F1. () Table of top-down regulated genes show highest fold change in EF-24–treated Kasumi-1 cells.

Pathway analysis further unveiled top canonical pathways, including the Pathogen Induced Cytokine Storm Signaling (PICSS) Pathway and IL-10 signaling, which were not only activated in EF-24–treated Kasumi-1 cells (Figures 6C) but also in HL-60 cells. Excessive production of inflammatory signals by the immune system can result in a cytokine storm, potentially leading to cell death.^34^ Moreover, TP53 and TP63, which function as tumor suppressors, appeared as upstream regulators of genes induced in EF-24–treated Kasumi-1 cells (Figure 6D).

Considering the downregulation of a substantial number of genes by EF-24, we investigated the pathways controlling these downregulated genes. Pathways crucial for cell cycle control, chromosomal replication, and DNA synthesis were downregulated, leading to a multifold decrease in the expression levels of core molecules such as *TOP2A*, *MCM7*, *MCM2*, *CDK1*, *MCM4*, and *MCM3* (Figures 6E, 6F). Additionally, TBX3 and E2F, identified as upstream regulators, were inhibited in EF-24–treated cells, resulting in the downregulation of their downstream target genes (Figure 6G, 6H, 6I).

### The antitumorigenic effects of EF-24 in THP-1 cells are mediated by interferon alpha/beta signaling

Our analysis of differential gene expression in the comprehensive transcriptomic data from EF-24–treated and untreated THP-1 cells has unveiled notable changes in specific genes (Figure 7A). Seeking a deeper understanding of the biological implications stemming from EF-24–induced gene dysregulation, we delved into IPA. Within the EF-24–treated THP-1 cells, we observed the activation of interferon alpha/beta signaling—a pathway acknowledged for its role in inhibiting malignant cell growth through programmed cell death (Figures 7B).^30^ This underscores that the antitumorigenic effects of EF-24 in THP-1 cells are mediated by immune signaling. The genes upregulated and regulated by interferon signaling play pivotal roles in various biological processes and molecular functions.

**Figure 7:**
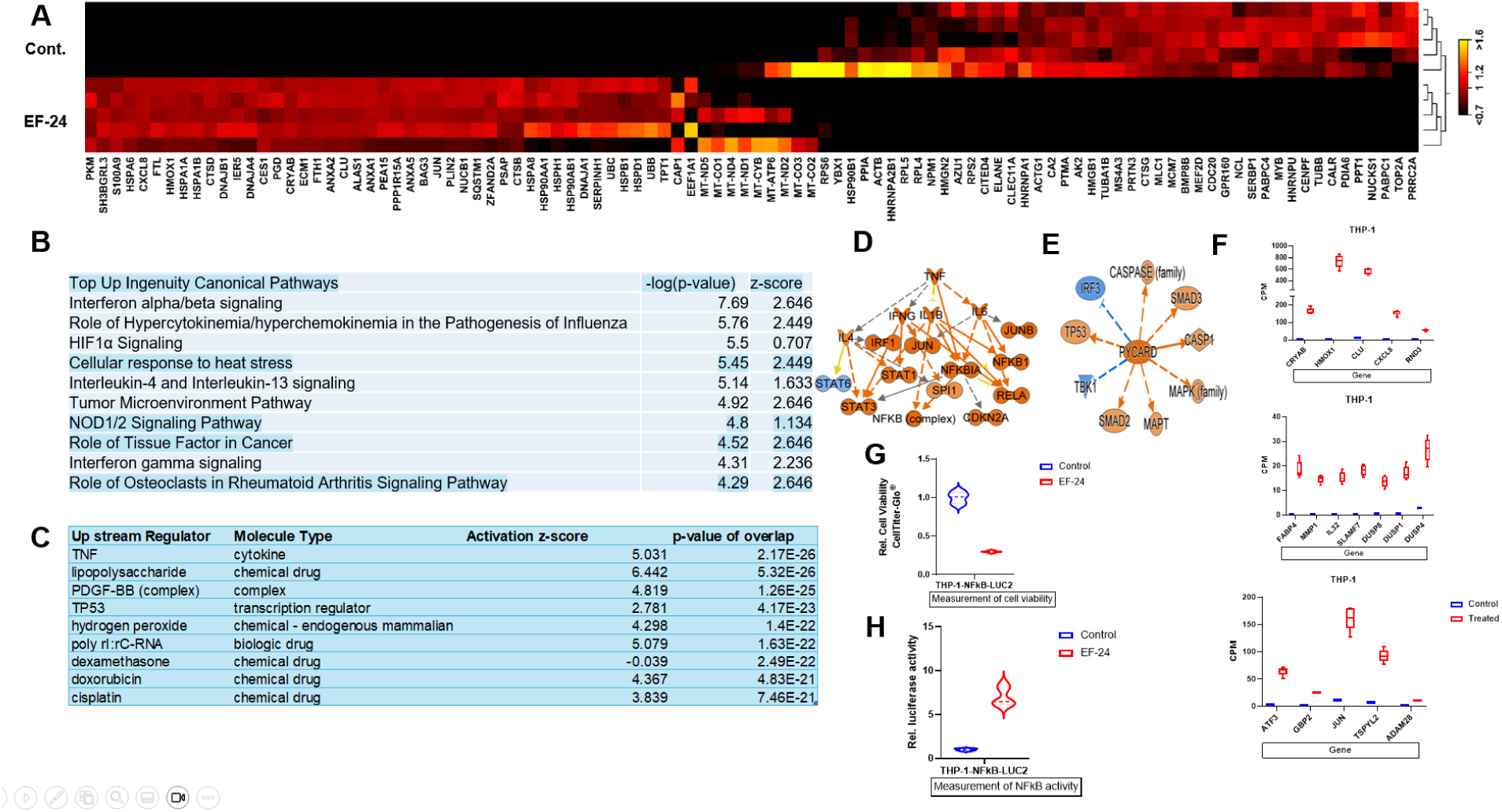
Differential gene expression pattern in THP-1 cells treated with EF-24. **(A)** Heatmap of the top 100 genes differentially expressed in EF-24–treated THP-1 cells (red = induced; black = reduced). **(B, C)** IPA revealed top canonical signaling pathways activated in EF-24–treated THP-1 cells. **(D)** TNF and **(E)** PYCARD activation observed in THP-1 cells treated with EF-24, functioning upstream of induced genes. **(F)** Box plots display quantitative levels of indicated genes in treated and untreated THP-1 cells. **(G, H,)** In EF-24–treated THP-1 cells, cell viability and NFκB activity show an inverse correlation. **(G)** Cell viability decreased in EF-24–treated THP-1 cells as compared to untreated control cells, whereas **(H)** NFκB-luciferase activity increased in EF-24–treated THP-1 cells as compared to untreated control cells.

Immunogenic cell death emerges as a significant mechanism in cancer therapy, harnessed by chemotherapy, radiation therapy, and targeted anticancer agents. This results in clinically relevant tumor-targeting immune responses^31^ and culminates in immunogenic cell death (ICD).^32^ In our exploration of upstream regulators influencing genes altered in EF-24-treated THP-1 cells, TNF and P53 emerged as key orchestrators of these changes (Figure 7C, D). TNF, acting as a death ligand, binds to its cognate receptors, instigating complex I and complex II, which regulate caspase-8 activation. While complex I is associated with cell survival and proliferation, complex II executes cell death by activating caspase-3 and caspase-7.^32^ P53, a master regulator, suppresses tumorigenesis and surfaced as a prominent upstream regulator in EF-24–treated THP-1 cells.^33^ P53 partially mediates the effects of EF-24 in inducing cell death in THP-1 cells.

Similarly, TNF regulatory networks displayed the activation of genes such as *IFNG*, *STAT1*, *JUN*, *CDKN2A*, *RALA*, and *NFKB1* in EF-24–treated THP-1 cells (Figure 7D). Activation of PYCARD resulted in downstream activation of genes such as *P53*, *MAPT*, *CASP1*, *SMAD2*, and *SMAD3* (Figure 7E). Subsequently, we identified candidate genes with differential expression and explored their biological associations; these genes exhibited a substantial increase in relative expression levels in EF-24–treated cells as compared to untreated THP-1 cells (Figure 7F). All these genes were found to be associated with cell death and survival, and their heightened activity contributed to induced cell death following EF-24 treatment.

To confirm TNF signaling activation in EF-24–treated cells, we performed a luciferase reporter assay using the THP-1-NFkB-LUC2 (ATCC^®^ TIB-202-NFkB-LUC2™) cell line, which was derived from parental THP-1 cells by introducing NFkB-LUC2 genes to measure NFkB activity. The cells were subjected to EF-24 treatment or left untreated for 24 hours, after which we measured cell viability and NFkB luciferase activity. As depicted in earlier assays, cell viability significantly decreased in the EF-24–treated cells as compared to untreated cells (Figure 7G). In contrast, NFkB activity saw a significant multifold increase (Figure 7H). This further substantiates the transcriptomics results and establishes the activation of TNF signaling in the EF-24–treated THP-1 cells.

## Discussion

In this study, we examined the impact of EF-24 on the transcriptome of leukemia cell lines.^34^ Despite numerous available treatments, myeloid leukemia – a disease with several genetically distinct subtypes – has seen few significant therapeutic advances.^35^ EF-24 is known to inhibit growth and induce apoptosis in leukemia and other cancers via established molecular pathways; however, its effects on global transcription remain unknown.^5,6,36–38^ To understand EF-24’s molecular mechanisms, we used next-generation sequencing to perform a transcriptome analysis on four well-established leukemia cell lines. Our goal was to identify gene expression signatures predictive of EF-24’s effects on cancer cells for future studies.

The genomic approach revealed that EF-24 has potent antitumor activity, significantly reducing cell viability across all tested lines. Transcriptomic analysis showed that EF-24’s effects are mediated by a combination of genes and signaling molecules rather than a single pathway, highlighting its complex mechanism of action. The activation of pathways such as the S100 family signaling pathway,^7^ along with the identification of key molecules like IL6 and TNF, provided nuanced insights into the intricate molecular dynamics influenced by EF-24.^8,39^ These findings underscore the agent’s multifaceted nature in modulating genes associated with cell survival and proliferation.

In our analysis of K-562 cells, we uncovered a myriad of differentially regulated genes post-EF-24 treatment, shedding light on its mechanisms. Surprisingly, EF-24 treatment induced NFkB activity in K-562 cells, contrasting with previous reports of NFkB suppression in EF-24 treated lung, breast, ovarian, and cervical cancer cells. ^40^ Additionally, unlike the inhibition of HIF1α observed in certain cell lines, HIF1α was activated in myeloid leukemia cell lines upon EF-24 treatment. ^41,42^ Accordingly, EF-24-mediated inhibition of NFkB and HIF1α in cancer cells appears to be contextual, and its antitumorigenic effects are not simply exerted by regulating specific genes or pathways. However, we observed that the increased expression of transporter genes ABCB1, ABCB11, ABCA4, ABCA9, and many others did not contribute to the antiproliferative effects of EF-24, which is consistent with previous reports by Skoupa et al. ^6^ Noteworthy inductions of apoptosis associated genes CLU and CRYAB, coupled with the activation of the wound healing signaling pathway unveiled EF-24’s potential in orchestrating cellular responses.^43,44^ This revelation emphasizes its prospective role in the context of chronic myeloid leukemia. Previous reports indicate that *Clusterin* (CLU) mediates apoptosis by interacting with Bcl-XL,^45^ aligning with the observed reduced cell viability in EF-24–treated cells. Attention was then directed to genes that exhibit higher levels of basal expression in untreated K-562 cells and overexpressed by EF-24, including *CLU*, *PTPRN*, *NDRG1*, *GBP2*, *OSGIN1*, *AFT3*, *IFI16*, *HLA-C*, *BEX2*, and *VWA5A*. Gene ontology analysis of the induced genes shed light on their potential antitumorigenic roles, suggesting tumor suppressor functions in EF-24–treated cells.^46–54^ Notably, activation of wound healing pathway suggests a counter response to the acute cell-killing activity of EF-24 treatment in K-562 cells. Induction of cell death is inexorably linked with cancer therapy, but this can also initiate wound-healing processes that have been linked to cancer progression and therapeutic resistance.^55,56^ Because the wound healing pathway was activated in EF-24–treated K-562 cells, understanding the therapeutic benefit of the curcumin analog EF-24 in chronic myeloid leukemia required further investigations.

Shifting our focus to HL-60 cells, EF-24 treatment induced significant alterations in the expression of genes associated with cancer progression. The downregulation of the neutrophil degranulation pathway suggested an unfavorable environment for leukemia cell survival, while the concomitant upregulation of IL-10 signaling hinted at a complex interplay between proinflammatory, and cell survival pathways triggered by EF-24.^57^ Examining gene regulatory networks associated with molecular and cellular function, we observed the activation of leukocytes and differentiation of hematopoietic progenitor cells in EF-24–treated cells. While the activation of leukocytes is known to be part of the immune response, the impact on tumor burden requires further exploration in an in vivo setting. Additionally, the activation of genes associated with the differentiation of hematopoietic progenitor cells suggests that EF-24 exerts multifaceted antitumor effects in HL-60 cells.^56^

Investigating upstream regulators of genes aberrantly activated in EF-24–treated cells, TNF emerged as a central regulator in HL-60 and THP-1 cells. TNF directly regulates upregulated genes, including *EGR1*, NFkB complex, *RELA*, and *IKBKB*, leading to downstream upregulation of *TP53*. EGR1 and TP53, known tumor suppressors, upregulate *p21*, inducing tumor cell apoptosis.^58,59^ Glucocorticoid receptor (NR3C1) signaling, activated by dexamethasone, a known treatment for leukemia, was found to be negligible changed, aligning with the observed negative control of genes regulated by dexamethasone in EF-24– treated cells HL-60 cells.^60^

In Kasumi-1 cells, EF-24 induced a distinct shift toward tumor-suppressive mechanisms. The activation of TP53 and TP63, coupled with the downregulation of tumor promoters, pointed towards EF-24’s potential to create a tumor-suppressive microenvironment. The modulation of key pathways, including the Pathogen Induced Cytokine Storm Signaling Pathway and IL-10 Signaling,^30,57^ added depth to our understanding of EF-24’s impact on gene regulatory networks.

Finally, our investigation extended to THP-1 cells, revealing EF-24’s prominent activation of the interferon alpha/beta signaling pathway.^30^ This immune-mediated pathway, coupled with the involvement of TNF and P53 as upstream regulators, substantiated EF-24’s potential to induce programmed cell death in malignant cells. Luciferase reporter assays further validated the activation of the TNF signaling cascade, reinforcing our transcriptomic findings.

## Conclusion

Our comprehensive study not only unraveled EF-24’s potent antitumorigenic activity and its influence on transcriptional landscapes but also delved into its multifaceted impact on specific signaling pathways and gene networks controlling cell survival, proliferation, and immune responses in myeloid leukemia cells. These insights lay a solid foundation for the exploration of EF-24 as a promising therapeutic agent in myeloid leukemia treatment. Moving forward, further investigations in preclinical and clinical settings are imperative to harness the full potential of EF-24 for effective leukemia interventions.

## Materials & Methods

### Cell Lines

Preserved cell vials of the cell lines K-562 (ATCC^®^ CCL-243™), HL-60 (ATCC^®^ CCL-240™), Kasumi-1 (ATCC^®^ CRL-2724™), and THP-1 (ATCC^®^ TIB-202™) were acquired from the ATCC^®^ repository (Manassas, VA, USA) and cultured according to specified parameters. These parameters were tailored for each cell line and adhered to in the ATCC^®^ cell culture laboratory, which operates under industry safety standard and quality guidelines and complies with ISO 9001 regulations. In brief, HL-60 cells were cultured in Iscove’s Modified Dulbecco’s Medium (IMDM; ATCC^®^ 30-2005™) supplemented with fetal bovine serum (FBS; ATCC^®^ 30-2020™) to a final concentration of 20%. Complete medium for K-562 consisted of IMDM with 10% FBS. For Kasumi-1, complete medium was prepared using RPMI-1640 (ATCC^®^ 30-2001™) supplemented with 20% FBS. THP-1 cells were cultured in RPMI-1640 supplemented with 10% FBS. All cell lines were maintained at 37°C in an atmosphere of 95% air and 5% CO_2_.

### Chemical and Reagents

EF-24 (with a purity of ≥98%; determined by HPLC (High Performance Liquid Chromatography)) was procured from Sigma-Aldrich^®^ (St. Louis, MO, USA; catalog number E8409-25MG). It was dissolved in dimethyl sulfoxide (DMSO; ATCC^®^ 4-X™) at a concentration of 25 mg (milligram) EF-24 per 2.5 mL (milliliter) DMSO, yielding a stock concentration of 32.1203 mM (millimolar) according to the manufacturer’s instructions. Subsequently, a 10 mM solution was prepared from the stock by combining 312.5 µL (microliter) of the 32 mM EF-24 solution with 687.5 µL of DMSO to achieve a final volume of 1 mL. The final treatment concentration was 2 µM; for instance, 10 µL of the 10 mM EF-24 solution was added to 50 mL of media to attain a final concentration of 2 µM in cell culture.

### Cytotoxicity Assay

The cytotoxic effect of EF-24 on AML cell lines was assessed using the CellTiter-Glo^®^ Luminescent Cell Viability Assay kit (Promega^®^ catalog number G7570). K-562, HL-60, Kasumi-1, and THP-1 were seeded into T25 flasks (Corning, part no. 430639) at a density of 3 x 10^6^ cells per flask. Subsequently, the cells were treated with a concentration of 2 µM EF-24 for 24 hours. Following treatment, 100 µL of cell suspension was combined with 100 µL of CellTiter-Glo^®^ solution in a 96-well plate with each well containing 3 x 10^4^ cells. The plate was then incubated at room temperature for 10 minutes, after which the absorbance was measured at 450 nm using a microplate reader (Molecular Devices SpectraMax^®^). Each experiment was performed twice with a minimum of five biological replicates.

### RNA Extraction and Quality Control (QC)

RNA isolation was performed using the QIAGEN^®^ QIAcube^®^ automated system with the RNeasy Mini QIAcube^®^ Kit. Frozen samples were thawed and prepared for RNA extraction according to ATCC’s work instructions. Extracted samples were tested for RNA integrity and quality using the Agilent^®^ TapeStation™ (RNA Integrity Number (RIN) ≥ 6.5), RNA purity using the Thermofisher^®^ Nanodrop™ (A260/A280 1.8 ≥ x ≤ 2.2), and concentration using the Qubit^®^.

### RNA-Seq Library Preparation and Sequencing

Automated RNA-seq NGS library preparation was performed on the Eppendorf epMotion^®^ 5075 Liquid Handler using the Illumina^®^ Stranded mRNA Prep, Ligation kit. Prepared NGS libraries were assessed by quantitative analysis using the Invitrogen^®^™ Qubit^®^™ dsDNA High Sensitivity Assay Kit and qualitative analysis using the Agilent^®^ 4200 TapeStation<u>^® TM^</u> and D5000 ScreenTape System. Libraries were loaded on an Illumina^®^ P3 200-cycle Reagent kit and sequenced on the NextSeq^®^ 2000 platform.

### RNA-Seq Data Analysis

Our data analysis pipeline underwent several meticulous steps, including quality control, read trimming, alignment to the reference transcriptome, and quantification of gene expression. Utilizing the QIAGEN CLC Genomics Workbench equipped with statistical and bioinformatics tools such as expression browser, principal component analysis (PCA), heatmap, correlation analysis, differential expression for RNA-Seq (CLCRNA-Seq Analysis Tools), and functional enrichment analysis, we ensured a robust analysis. The detailed workflow of the experiment and data processing pipeline is presented in Supplemental Figure 1 (CLC workflow). Adhering to stringent quality criteria—with total minimum reads exceeding 18M and genome coverage surpassing 85%—was crucial for the integrity of our analysis.

### Bioinformatics Analysis

The gene list obtained from the differential gene expression analysis underwent Ingenuity Pathway Analysis (IPA). IPA generated comprehensive reports on the top canonical pathways, upstream regulators, diseases and biofunctions, regulator effect networks, networks, tox lists, machine learning (ML) disease pathways, and analysis-ready molecules.

### Statistical Analysis

Values are presented as the mean ± SD (Standard Deviation). Statistical analyses were conducted using the Microsoft excel student’s t-test and GraphPad Prism^®^ software.

## Data Availability

All RNAseq data produced for this study is available under NCBI Bioproject PRJNA1165739.

## Authors Contributions

Conceptualized the project, A.P.S. and J.L.J.; Designed the methodology, performed experiments, analyzed data, lead, and supervised all aspects of the project, wrote the manuscript, A.P.S.; Analyzed data, review and edit the manuscript, supervised the project, J.L.J; RNA extraction and sequencing; N.W., J.D., A.F. All authors have read and agreed to the published version of the manuscript.

## Acknowledgments

We thank Cara Wilder for helping edit and review the manuscript.

